# Experimental disruption of social structure reveals totipotency in the orchid bee, *Euglossa dilemma*

**DOI:** 10.1101/2022.01.27.478072

**Authors:** Nicholas W. Saleh, Jonas Henske, Santiago R. Ramírez

## Abstract

Eusociality has evolved multiple times across the insect phylogeny. Social insects with greater levels of social complexity tend to exhibit specialized castes with low levels of individual phenotypic plasticity. In contrast, species with small, simple social groups may consist of totipotent individuals that can transition among behavioral and reproductive states as the social hierarchy shifts. However, recent work has shown that in some simple social groups, there can still be constraint on individual plasticity, caused by differences in maternal nourishment or initial social interaction. It is not well understood how and when these constraints arise during social evolution, ultimately leading to the evolution of nonreproductive workers. Some species of orchid bees can form social groups of a dominant and 1-2 subordinate helpers where all individuals are reproductive. Females can also disperse on emergence to start their own nest as a solitary foundress, which includes a nonreproductive nest guarding phase not typically expressed by subordinates. Little data exist to characterize the flexibility of orchid bees across these trajectories. Here, using the orchid bee *Euglossa dilemma*, we conduct an experiment assessing the plasticity of subordinate helpers, finding that they are highly flexible and capable of the behavioral, physiological, transcriptomic, and chemical changes seen in foundresses. Furthermore, we identify genes and gene networks associated with reproductive changes in *E. dilemma* that overlap with genes associated with worker physiology in eusocial species. Our results provide evidence that the lack of nonreproductive workers in *E. dilemma* is not due to a lack of subordinate plasticity.

## Introduction

The evolution of obligate eusociality, such as seen in ants, honey bees and termites, is expected to result in a transition of plasticity from the individual level to the colony level (Taylor et al., 2019). Species with obligate eusocial behavior may exhibit irreversible castes with morphological traits that are adapted to specific tasks within the colony (*e.g*. soldiers), with these traits being determined during development (Rehan and Toth, 2015). In contrast, individuals of species forming small, cooperatively breeding groups are often totipotent as adults, with any member of the social group exhibiting the flexibility to serve as the primary reproductive, with dominant and subordinate roles defined after eclosion (Johnson and Linksvayer, 2010; Strassmann et al., 2002). While there is substantial debate about the life-history features present in the ancestors of obligate eusocial species (Linksvayer and Johnson, 2019), extant species with small social groups are frequently used as model systems to evaluate hypotheses about solitary to social life-history transitions (Kronauer and Libbrecht, 2018).

Empirical study has shown that, while these species do show higher adult flexibility than obligately eusocial species, they may still experience some constraints on their adult plasticity, either through alternative developmental trajectories or social interactions (Lawson et al., 2017; Awde and Richards, 2018). Understanding how and when changes in plasticity first arise is important in identifying the mechanisms leading to the evolution of fixed, nonreproductive worker castes that are developmentally determined (Linksvayer et al., 2011; Jones et al, 2017). In the small carpenter bee *Ceratina calcarata*, for instance, mothers may undernourish their first female offspring, creating a small-bodied helper which does not reproduce on their own but provisions their siblings, which will disperse to start their own nests (Lawson et al., 2016). These helper individuals have been suggested to represent “caste-antecedents,” showing extensive overlap in gene expression patterns with eusocial workers (Shell and Rehan, 2019). Similarly, the facultatively eusocial Halictid bee *Megalopta genalis* appears to rely on maternal manipulation of offspring provisions to create small-bodied females that become nonreproductive workers (Kapheim et al., 2011), though these workers can still assume a vacant queen position and reactivate their ovaries if given the opportunity (Jones et al., 2017). In contrast, adults of some species forming small social groups show no apparent signs of constraint on adult plasticity. Some primitively eusocial hover wasps and some allodapine bees, for example, have little to no consistent body size differences that correlate with social hierarchy, with all adults capable of any social or reproductive role (Field et al., 1999; Sumner et al., 2002; Schwarz and Woods, 1994). Ultimately, to understand how the specific life-history features of species with simple social groups relate to the evolution of eusociality requires evaluation of these features in a phylogenetic context (Linksvayer and Johnson, 2019; Shell et al., 2021).

Uncovering the evolutionary history of sociality in the corbiculate bees (honey bees, bumblebees, stingless bees, and orchid bees), most of which are well-known for their complex obligately eusocial colonies, has been hampered both by phylogenetic uncertainty regarding the relationships among lineages (Engel and Rasmussen, 2020) and by the apparent lack of closely related species showing small or intermediate social group sizes (Danforth, 2002). While recent work has reduced much of the phylogenetic uncertainty among the corbiculate bee lineages (Romiguier et al., 2016; Bossert et al., 2017), the lack of data to inform the seemingly abrupt evolution of eusocial behavior remains a challenge. However, part of this difficulty arises due to the lack of information about life-history variation among the orchid bees, the earliest branching lineage of the corbiculate bees, which have primarily been considered to be solitary (Cameron, 2004; Fischman et al., 2017).

Indeed, the state of orchid bee social behavior has been a puzzlement for biologists, which have, until somewhat recently, relied on rare observations of nesting to characterize behavior across the 200+ orchid bee species (O’Toole and Raw, 1991; Ramirez et al. 2002). After observing that some orchid bees had multiple preconditions favoring eusociality, such as overlapping generations, long lived individuals, and semipermanent nests, for example, Roberts and Dodson (1967) posed the question, “why, then, has there been no evolution of distinct worker and reproductive castes among these bees?” As new data emerge, however, it is increasingly clear that numerous orchid bee species do show diverse social behaviors, though there is still no evidence for true nonreproductive castes among individuals in a social group, which have always been found to have activated ovaries. Social behaviors documented among orchid bee species include communal nesting, multi-female nest founding, overlapping generations, and the division of labor between dominant nest guards and subordinate foragers (Augusto and Garófalo, 2004; Capaldi et al., 2007; Cocom Pech et al., 2008; Solano-Brenes et al., 2018).

In the orchid bee *Euglossa dilemma*, the focal species of this study, nests are started by a single foundress that constructs a nest of plant resin and provisions an initial brood batch with pollen and nectar. After completing these brood cells, the foundress ceases foraging and reproduction and transitions into a “guard” phase to protect her developing brood. When a foundress enters this nonreproductive guard phase, her ovaries inactivate and reduce in size. This shift to guard behavior is associated with numerous changes in gene expression across the brains and the ovaries, including genes associated with social behavior in eusocial species (Saleh and Ramírez, 2019). After spending up to two months in the guard phase, offspring emerge, and the nest enters the social phase. During this transition to social behavior, the foundresses’ ovaries reactivate and she then becomes the dominant bee, while 1-2 of her female offspring may remain in the nest as a subordinate helper. Other female offspring disperse to begin their own nests. Between individuals in a social nest, there is division of labor, with the dominant bee remaining in the nest with the brood while the subordinate bee forages for new offspring. Both the dominant and subordinate bee are reproductive; however, the dominant bee eats and replaces all subordinate laid eggs, indirectly resulting in a functional reproductive division of labor.

Like several other orchid bee species, *E. dilemma* shows behavioral plasticity among social roles. Subordinates can transition from a subordinate to dominant position when the dominant is removed in a multifemale nest (Andrade-Silva and Nascimento, 2016; Séguret et al., 2021; N. Saleh, personal observation). Although this plasticity is notable, the transition is between two reproductive behaviors that show relatively slight physiological differences (Saleh and Ramírez, 2019). In contrast, the transition from the foundress phase (reproductive) to the guard phase (nonreproductive) in the solitary portion of the lifecycle is pronounced and involves substantial behavioral and physiological changes (Saleh and Ramírez, 2019). However, it is unclear if subordinates, which remain in their natal nest as foragers, can express the full range of plasticity shown by dispersing foundresses. This is unclear because the guard phase, which occurs after the provisioning of the first brood, can be entirely absent in orchid bee social nests, due to continuous generations as nest size grows (Augusto and Garófalo, 2009; Boff et al., 2017). Alternatively, when a social nest does show an interval of reproductive inactivity between broods, it may be relatively short, or the subordinate bee may abandon the nest early (Augusto and Garófalo, 2011). Consequently, this plasticity is highly variable in subordinate helpers relative to the predictable, prolonged changes seen in solitary foundresses.

In this study, we seek to assess the behavioral and reproductive plasticity of the social orchid bee *Euglossa dilemma*, testing the hypothesis that subordinate helpers in *E. dilemma* social groups can regulate their reproductive physiology dynamically, expressing both reproductive and nonreproductive phenotypes, despite these nonreproductive phenotypes being absent in typical social interactions. This study aims to provide insight into whether the lack of true reproductive castes in *E. dilemma* is, in part, due to a lack of reproductive plasticity in subordinates. We assess this by isolating individual subordinates and disrupting their social behavior to simulate conditions experienced by solitary foundresses starting their own nest. We then collect behavioral, physiological, chemical, and transcriptomic data from these isolated subordinates to determine the degree to which phenotypic changes mirror those of solitary foundresses.

## Methods

### Nest observation

All nest observations were conducted in Ft. Lauderdale, FL, where a naturalized *E. dilemma* population has been present for around 15-20 years (Skov and Wiley, 2005). Wooden nest boxes were placed on the eaves of buildings in Ft. Lauderdale in which *E. dilemma* females naturally founded nests. Transparent red plexiglass lids were placed on top of these wooden boxes to facilitate video recording and behavioral observation. In some nests, infrared CCTV cameras were used to record 24hr continuous video through the lid on top of the nest boxes. We also surveyed nests daily, checking nest occupancy. In the evening, following the return of all bees to the nest, individual bees were tagged with small numbered, plastic discs superglued to the thorax.

### Nest manipulation

To test the hypothesis that subordinate helpers will express the plasticity exhibited by foundresses, we first identified nests containing subordinate individuals and then we experimentally remove the interaction with other dominant or subordinate nestmates to determine how their behavior progresses in isolation. Our approach is illustrated in Fig. 1 and possible outcomes are illustrated in Fig. S1. First, in the summers of 2018 and 2019, we identified nests in the guard phase, where offspring from the first brood had not yet emerged. Following offspring emergence, we waited until individuals remaining in the nest showed consistent dominant/subordinate relationships before manipulation. We define an individual as subordinate if it has provisioned at least one brood cell with subsequent egg replacement by the dominant (the original female in the nest, which in most cases is the mother). After this is confirmed, we removed all individuals from the nest except the first bee that showed consistent subordinate behavior. Removal of individuals occurred after dark, to confirm that all bees had returned to the nest after foraging. The subordinate bee left behind was not handled in this process. The time elapsed before nestmate removal varied among nests, to ensure that all offspring in the brood cells had emerged. If additional females emerged from brood cells after nestmate removal, the remaining subordinate could transition to dominant behavior and there would be no opportunity for that subordinate to express the nonreproductive changes seen in foundresses. In three of 14 nests where this manipulation was performed, several offspring from the first generation failed to emerge (due to disease, parasitism, or unknown causes) and these brood cells were carefully cut from the nest using a sterilized razor when nestmate removal occurred.

**Figure 1.**
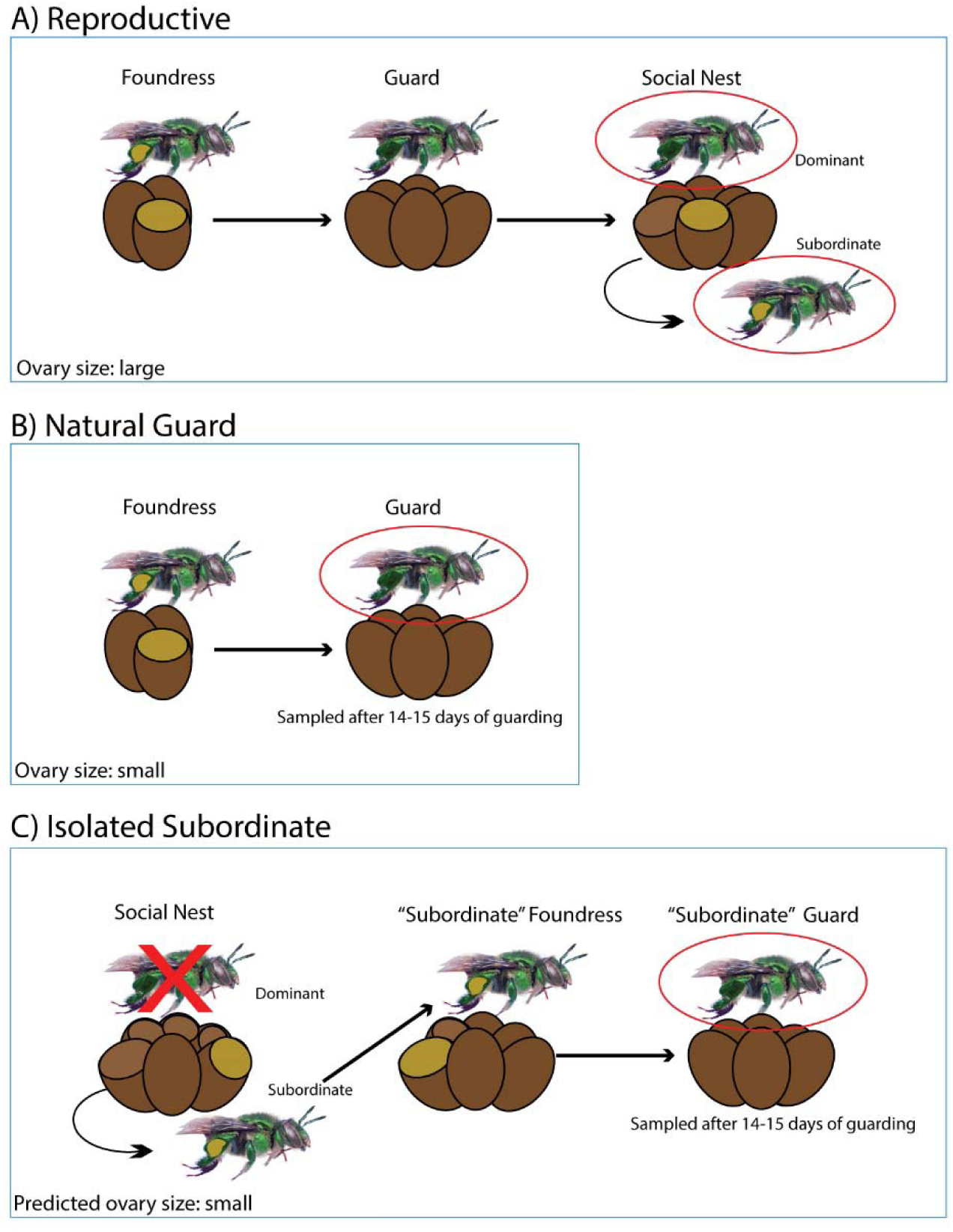
Design for subordinate isolation experiment, illustrating behavioral progression of individuals and nests from the three different sampled groups, A-C. The blue boxes encompass the entire behavioral sequence of each group and red ellipses show which specific behaviors were sampled from these groups. The ovary size or predicted ovary size of the sampled individuals are listed in the blue boxes. Yellow on the hindleg (corbicula) or in the brood cell represents pollen and ongoing provisioning. A light brown ellipse on top of a brood cell indicates that offspring have emerged and that the brood cell is empty. **A)** Natural nest progression, where individuals are sampled performing reproductive behaviors (dominant and subordinate). **B)** Naturally guarding individuals sampled 14-15 days after showing guard behavior. **C)** Isolation treatment where the dominant individual was removed (indicated by the red “X”). Isolated subordinates that transitioned to guarding behavior were collected after 14-15 days.

After nestmate removal was conducted, we observed the behavior of the remaining subordinate to monitor changes in foraging behavior and/or the start of guarding behavior. We classify an individual as in the “guard” phase if it has discontinued all pollen foraging trips and remains inside the nest with a resin seal over the nest entrance during normal foraging hours (sunrise to sunset) on a day where foraging is seen in other nests. Individuals that successfully became guards were collected after showing 14-15 days of consistent guarding behavior. In total we performed 14 removals. Individuals collected from these treatments are hereafter referred to as “isolated subordinates.”

### Guard and reproductive individuals for comparison

To compare the changes in isolated subordinates to those occurring naturally in dispersing foundresses, we observed and collected a set of control individuals, which constructed nests as solitary foundresses before they naturally transitioned to guarding behavior, for 14-15 days (n =9). We hereafter refer to these individuals as “natural guards.” We also recorded brood size for several additional natural guard individuals not collected or disturbed (n = 4). In addition, we collected dominant (n =5) and subordinate (n = 5) individuals, to compare reproductive phenotypes (dominants and subordinates) to nonreproductive phenotypes (isolated subordinates and natural guards). This allows us to assess whether isolated subordinate phenotypes more closely resemble undisturbed bees at the reproductive stage (dominant and subordinate) or undisturbed bees at the guard stage.

### Sample Collection

To collect individuals after observation, entire nest boxes were placed on dry ice to incapacitate bees, which were then removed from the nest and immediately frozen in liquid nitrogen for subsequent RNA extraction and sequencing. The entire collection process was completed within minutes of nest box removal from the field. Collection of all individuals occurred between 12-4 pm during normal afternoon foraging. After storage in liquid nitrogen for 1-3 weeks, samples were transferred to a -80°C freezer until further phenotypic analysis. Three isolated subordinates were collected in the Summer of 2018, with all other isolated subordinates, natural guards, and dominants and subordinates collected in the Summer of 2019. An extreme weather event resulted in a truncated collection season in the Summer of 2019, requiring collection of all dominant and subordinate samples as well as one isolated subordinate sample on a single day. This isolated subordinate individual was collected after 10 days of guarding, in contrast to all other natural guards and isolated subordinates, which were collected 14-15 days after the start of guarding behavior. We assess the possible impact of these collection irregularities in appendix 1 found in the supplemental material.

### Ovary size measurement

An ovary size index was calculated using the sum of the longest basal oocyte in each ovary (two measurements), divided by the intertegular distance, to account for body size differences. We refer to this measurement when “ovary size” is mentioned. Oocyte length and intertegular distance were measured without knowledge of treatment/behavior to avoid possible bias in measurement. We also compare individuals in this study to *E. dilemma* guarding individuals from Saleh and Ramírez, 2019. The ovary size index from individuals in Saleh and Ramírez, 2019 is available only with measurements from the longest basal oocyte (as compared with the longest basal oocyte of each ovary, measured in this study). Consequently, we adjust our ovary size index to this slightly different approach only when comparing samples between the two studies.

### General statistical analysis

All statistical analysis was conducted in R version 3.6 (R Core Team, 2020). For assessing differences among mean values (such as between ovary size index values, body size measurements, or brood sizes), one-way ANOVAs were used, with Tukey’s HSD tests to assess pairwise relationships among groups. We used a Levene’s test and a Shapiro-Wilk test to verify ANOVA assumptions. If either assumption was violated, we proceeded instead with Kruskal-Wallis tests using Steel-Dwass tests for pairwise comparisons (Douglas & Michael, 1991). All statistical tests were done with reproductive individuals (subordinates and dominants) considered together as one group.

### CHC extraction, data generation, and analysis

Cuticular hydrocarbon differences, which are associated with behavior in *E. dilemma* (Saleh et al., 2021), were extracted from one pair of fore and hindwings, as in Martin et al., 2009, by placing them in 100 μl of hexane for 10 minutes, occasionally swirling them. After 10 minutes, hexane was transferred to a GCMS vial and left overnight in a fume hood to evaporate. The next day, 30 μl of hexane was transferred to the vials, which were then run on the GCMS. Wing extracts have been shown to accurately reflect CHCs on the abdominal surface of *E. dilemma* females (Saleh et al., 2021).

Sample extracts were run using a 1μl splitless injection on a GC-MS (Agilent 7890B GC, 5977A MS), with modifications to the protocol from Choe et al, 2012, which started at 100°C for 1 minute, increasing 15°C per minute until 300°C was reached, after which the program held at 300°C for three minutes. Helium was used as the carrier gas.

Chromatograms from the GC-MS were integrated to include peaks with an area corresponding to at least 0.1% of the largest peak. Chemical identification was accomplished by comparing to available data for *E. dilemma* (Pokorny et al., 2014; Pokorny et al., 2015, Saleh et al., 2021). and to mass spectral libraries and known mass indices. We excluded peaks that were not identified as CHCs (linear and branched alkenes and alkanes). After we removed the non-CHC peaks, we calculated the relative abundance of each CHC peak per sample by summing the total area of all peaks for that sample and then dividing the area of each individual peak by the total, generating proportional data. In addition to the individuals used in the rest of the study, CHCs from several additional dominants (additional n = 3, total n = 8) and subordinates (additional n = 3, total n =8) collected from the same field seasons, were available and included in the analysis.

For visualizing CHC differences, we used an NMDS plot based on Bray-Curtis dissimilarity among samples, as implemented in the Vegan R package (Oksanen et al, 2019). *Euglossa dilemma* has a well characterized CHC polymorphism segregating in Florida populations that complicates chemical comparison among samples but does not appear to be related social behavior (Saleh et al., 2021). Because of this, we exclude samples from the rarer CHC morph (n = 4) from NMDS analysis and statistical analysis for clarity (remaining n = 31). We show the same plot with all samples in supplemental figure S2 (n = 35) and data from all individuals is included in the supplementary data. We perform PERMANOVA analysis on the set of 31 individuals with 1000 permutations to assess the statistical significance of differences in CHC profile among groups. We used the RVAidememoire R package (Hervé, 2021) to perform contrasts with FDR correction for multiple comparisons.

### RNA extractions, sequencing, and quality control

For brain dissections we first removed the cuticle around the frons and post-occiput while samples were on dry ice. Next, frozen heads with the cuticle removed were placed in RNAlater ICE for at least 16 hrs at -20°C. After RNAlater ICE thaw, brains were dissected from the heads on dry ice and immediately transferred to Trizol solution for RNA extraction. We removed the retinas from the optic lobe as well as the ocelli during dissection, but otherwise the whole brain was included. We dissected the ovaries by first removing sections of abdominal cuticle from frozen samples on dry ice. The abdomens were then thawed in RNAlater ICE for at least 16 hrs at -20°C before being dissected on dry ice. Ovaries were photographed with a scale bar for measurement and then immediately placed in Trizol solution. We followed the standard RNA extraction Trizol protocol, with glycogen added to the brain samples but not the ovary samples to help increase yield. After extraction, RNA was cleaned using an Invitrogen Turbo DNA-free kit and then quantified using a Qubit. Next, RNA quality was checked on a Bioanalyzer (Agilent) and library construction commenced on samples with high quality RNA. These samples consisted of 26 brains (dominant = 5, subordinate = 5, natural guard = 7, isolated subordinate = 9) and 29 ovaries (dominant = 5, subordinate = 5, natural guard = 9, isolated subordinate = 10).

RNA samples were submitted for library preparation and sequencing to the Vincent J. Coates Genomic Sequencing Laboratory at UC Berkeley. Libraries consisting of 150 bp paired end reads were sequenced on a Novaseq 6000, generating an average of 30 million reads per library (mean = 30.40 million, S.D. = 4.89 million, range = 23.47 - 47.90 million, N = 55). After sequencing, we evaluated the quality of reads with FastQC (version 0.11.7, Andrews, 2010). Initial sample clustering using MDS in EdgeR suggested one brain sample from a natural guard individual to be an outlier with no obvious biological explanation. Furthermore, inspection of the FastQC reports for this individual showed some quality score drops in sequencing quality not shown in the other samples, so this individual (SRNS33) was dropped from all gene expression analysis (Figure S3). Two other brain samples clustered separately from the other samples along one axis in the MDS plot of gene expression data (SRNS11 and SRNS53; Fig S3); however, these were a dominant and subordinate gathered from the same nest. No technical reason could be identified that drove this pattern (they were collected at the same day/time as other samples and showed no obvious sequencing anomalies). Consequently, it appeared most likely that biological variation associated with shared nesting may be responsible for this pattern and so samples were included in our analysis, which aims to capture realistic levels of biological variation found in field established nests. The raw data for all sequenced samples can be found at NCBI under the bioproject accession PRJNA750777.

### Differential Gene Expression Analysis

We followed the same analytical approach as presented in Saleh and Ramírez, 2019, to facilitate comparison of results (as shown in appendix 1). Briefly, we used Kallisto (Bray et al., 2016) for producing transcript counts based on genes from the *E. dilemma* genome (Brand et al., 2017). After transcript quantification, we filtered genes in the ovary and brain data set separately, so that each of the two data sets consisted of genes with at least one count per million (CPM) in at least 5 of the libraries, which represented the smallest behavioral group sample size. For the brain data, this resulted in 11,041 genes and 10,132 genes for the ovary data filtered down from the total gene set of 16,127 genes. We used edgeR-robust (Zhou et al., 2014) with default settings and the glmLRT function with FDR <0.05 to identify differentially expressed genes (DEGs) among the four sampled behavioral groups. We used TMM normalization to account for differences in the amount of reads among libraries. Hierarchical clustering and heatmap construction was conducted using Euclidean distance and Ward.D2 clustering using gplots version 3.0.1 (Warnes et al., 2020).

### Gene network analysis

We conducted gene network analysis to identify co-expressed networks of genes underlying ovary size differences among individuals. This analysis may provide additional insight into functional connections between sets of genes underlying our phenotypes of interest that may not be captured during standard differential expression analysis (Faragalla et al., 2018). To do this, we used the WGCNA (weighted gene co-expression network analysis) package in R (Langfelder & Horvath, 2008) to identify modules of genes showing co-expression. We then used the module eigengenes, which summarize the expression of each module, to assess correlation between ovary size index measures and gene expression. WGCNA parameters followed recommended values from published tutorials and R code used for the analysis can be found in the supplemental data. Briefly, using the filtered and normalized gene sets from differential expression analysis, gene modules were detected using a soft thresholding power for which the scale free topology index value was greater than 0.85. For ovaries this soft thresholding power was four and for the brains it was five. The minimum module size was 30 genes and the module merging cut-off value was 0.25.

### Gene list comparisons

We compared genes identified through differential expression analysis and WGCNA to genes previously identified as associated with reproductive plasticity in other bee species. Specifically, we compared the results of this study to data from the abdomens of *Megalopta genalis* queens and workers (Jones et al., 2017) and the abdomens of *Apis mellifera* egg laying or non-egg laying workers (Galbraith et al., 2016). These comparisons represent two origins of eusociality and different levels of eusocial organization. *Apis mellifera* has a complex eusocial organization with colonies consisting of thousands of individuals but, phylogenetically, it is more closely related to *E. dilemma* than *M. genalis* and thus may share features associated with behavior in the mostly eusocial corbiculate bees. In contrast, *M. genalis* forms small, facultatively eusocial groups that more closely resemble the social structure of *E. dilemma*, though social behavior has arisen independently in these groups. In addition, we compared our results to genes identified as differentially expressed between natural guards and subordinates in Saleh and Ramirez, 2019, to assess the degree to which our results overlap. For comparisons to *M. genalis*, we identified orthologous genes between the published *E. dilemma* peptide set and the predicted peptides from Jones et al, 2017 using a reciprocal best hit (RBH) blastp search (e-value <1E-5). For *A. mellifera* comparisons, we converted our *E. dilemma* gene lists into honey bee gene IDs (OGSv 3.2; Elsik et al., 2014) with a conversion list from Brand et al, 2017. We used DAVID 6.8 to perform GO term analysis with Benjamini-Hochberg corrected p-values using honey bee OGSv 3.2 gene IDs. To identify significant overlaps between any two compared gene sets, we performed hypergeometric tests to identify overlaps greater than expected by chance when compared to the shared universe of analyzed orthologous genes.

## Results

### Behavioral response of isolated subordinates

We conducted 14 nest manipulations on naturally colonized nest boxes in the field, removing all individuals except for a single subordinate bee (Table S1, Table S3). In these 14 manipulated nests, three isolated subordinates disappeared before transitioning to guarding behavior. Two of these three disappeared after completing one additional brood cell following isolation and one disappeared the morning after isolation. One isolated subordinate disappeared from the nest after guarding began, but before collection. The remaining 10 bees successfully transitioned to guarding behavior and were collected and processed for subsequent analysis. Four of these 10 females did not provision additional brood cells after isolation before transitioning to guard behavior, ceasing foraging and brood cell construction upon isolation. Six of 10 females provisioned at least one brood cell after isolation before guarding behavior began. We also compared the final brood size of isolated subordinates and naturally guarding individuals, finding that isolated subordinates began guarding a smaller number of brood cells on average (mean = 5.3, S.D. = 1.8, range = 3-9, n =11;) compared to naturally guarding bees (mean = 7.8, S.D. = 2.4, range = 5-14, n =13). These differences are significant, though there is a substantial overlap between them (F_1,22_ = 7.62, p = 0.011, Fig. S4).

### Isolated subordinates exhibit reduction in ovary size

We examined the ovary size of isolated subordinates collected after showing guarding behavior, comparing them to naturally guarding bees and reproductive individuals (dominants and subordinates). We find that isolated subordinates and natural guards show a reduction in ovary size relative to reproductive individuals, though ovaries of isolated subordinates and natural guards are statistically indistinguishable from each other (F_2,26_ = 33.08, p < 0.001; Fig. 3). We find no difference in body size among these three groups (Kruskal-Wallis χ^2^ = 2.7, df = 2, p = 0.26, Fig. S5). Given the lack of ovary size index differences between the isolated subordinates and natural guards, we combined these groups (n =19) and compared their ovary size index measurements to ovary size index measurements from naturally guarding individuals measured in Saleh and Ramírez, 2019 (n = 15). The individuals in Saleh and Ramírez, 2019 were collected after performing guarding behavior for a longer time range (minimum two weeks with some samples likely up to six weeks). Consequently, comparison to those samples can indicate whether the reproductive transition measured in this study is complete or if the reduction in ovary size would continue beyond two weeks into the guarding phase. We find that our sampled individuals, which guarded for 14-15 days, had larger ovaries on average than natural guards from Saleh and Ramírez, 2019 which guarded for longer periods on average (F_1,32_ = 10.25, p < 0.01; Fig. S6).

**Figure 2.**
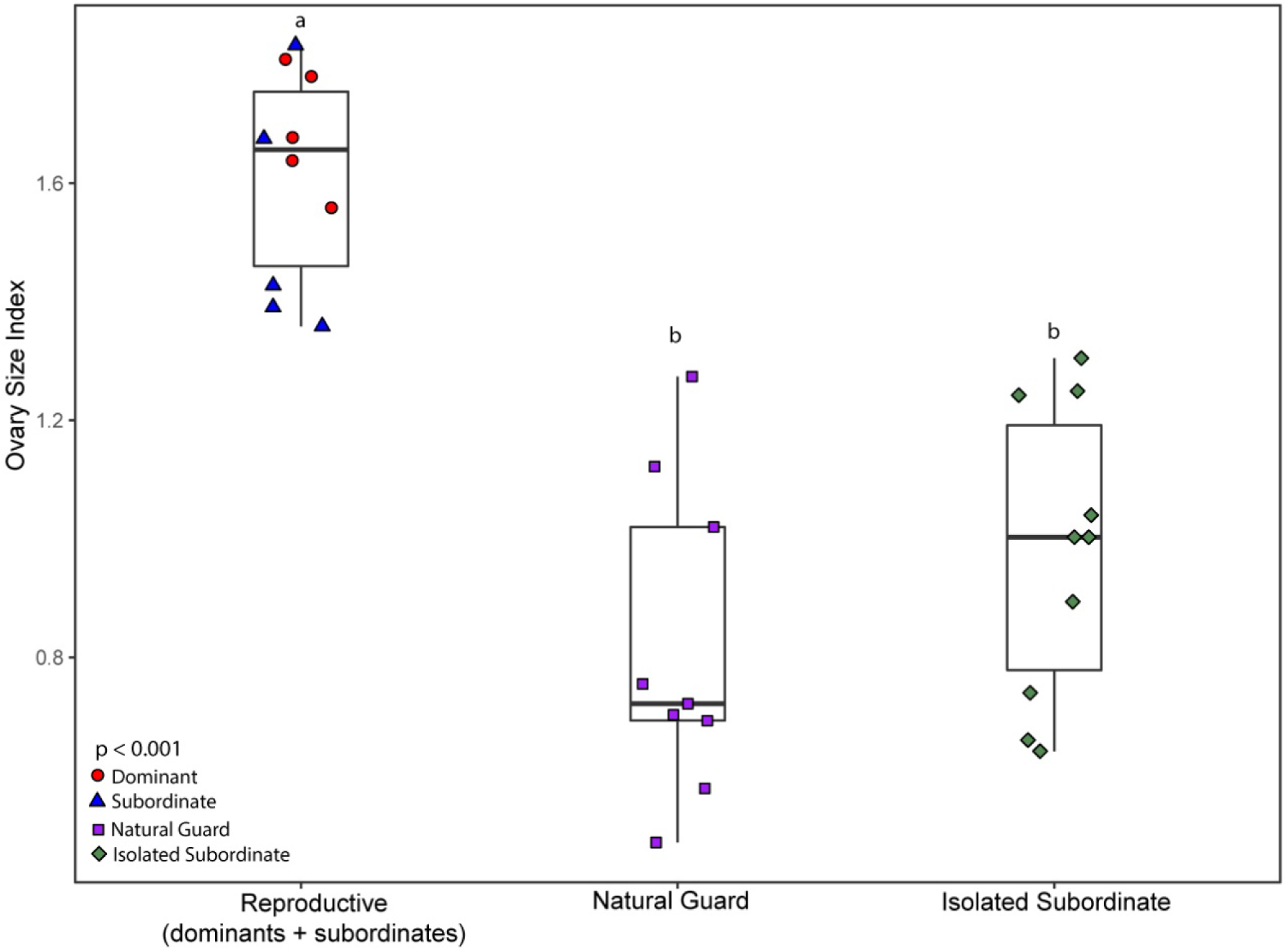
Ovary size index among reproductive individuals, natural guard individuals, and isolated subordinate individuals. Letters indicate statistical groupings determined by a Tukey HSD test with the p-value calculated using a one-way ANOVA. The box plots show the mean value in each group.

**Figure 3.**
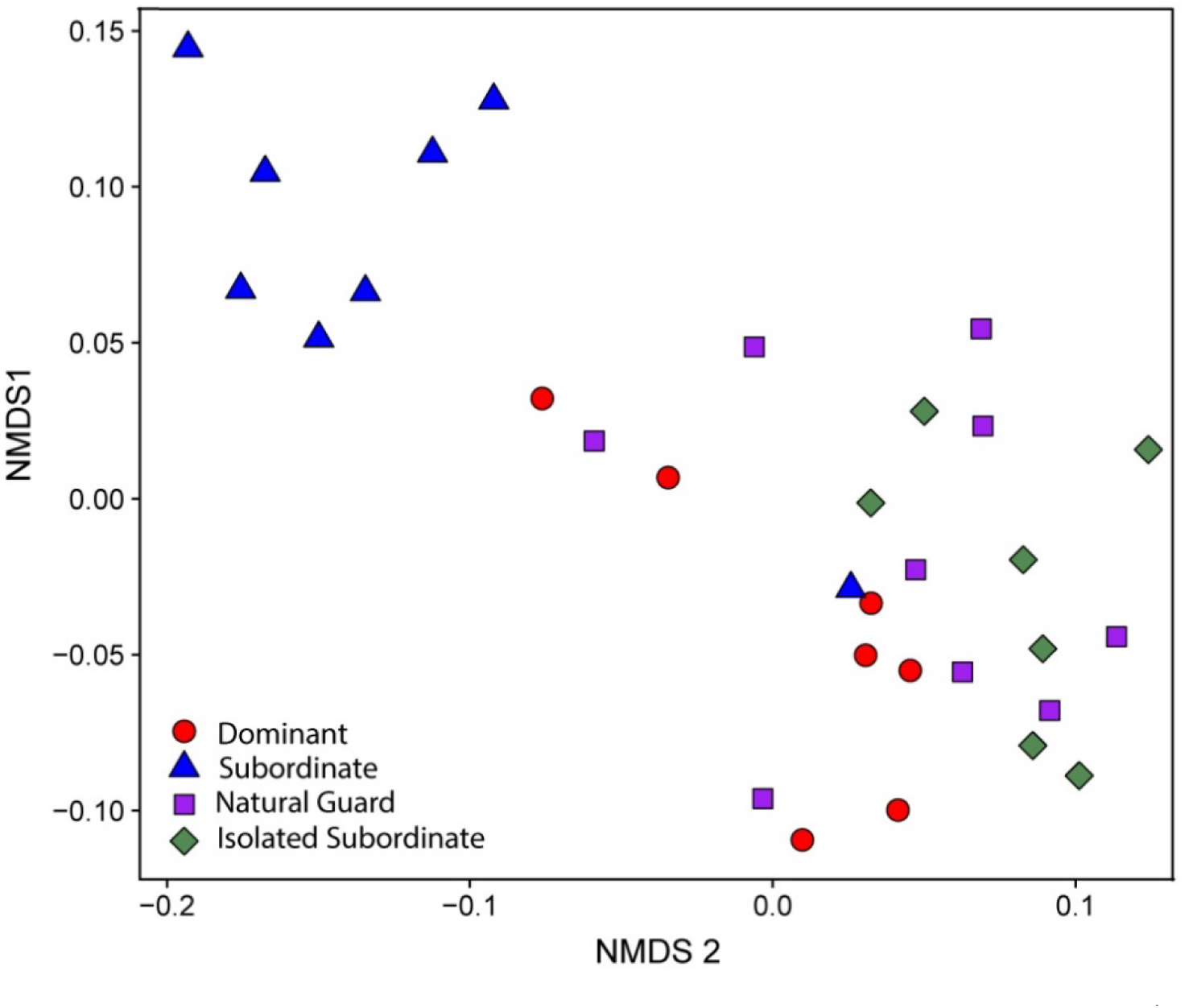
NMDS plot of CHC variation across behaviors. Unique color/symbol combinations represent the different behavioral groups. Stress value for NMDS configuration = 0.032.

### Isolated subordinates show shift in CHCs

We examined variation in the CHC profiles, finding 17 previously characterized alkanes and alkenes (Saleh et al., 2021), and identified changes that may be associated with behavior and reproduction. In contrast to the ovary size data, individuals do not separate strictly based on reproductive state, with dominants, isolated subordinates, and natural guards mostly clustering separately from subordinates (Fig. 3). Consistent with this, pairwise PERMANOVA with FDR adjustment finds subordinates significantly differentiated from all three other groups, primarily driven by an increased relative abundance of shorter chain alkanes between 21 and 24 carbons long. Isolated subordinates and guards are not statistically distinguishable from each other (p = 0.29). Dominants and natural guards are not statistically distinguishable from each other (p = 0.29), though isolated subordinates and dominants are significantly different (p = 0.013).

### Isolated subordinates show gene expression patterns consistent with guarding behavior

Our gene expression analysis had two aims: (1) identify gene expression patterns associated with the sampled behaviors and (2) determine whether isolated subordinates resemble natural guards or other behavioral phases based on their expression profiles. To this end, we first found all DEGs among pairwise comparisons of the four sampled behavioral groups in the EdgeR model. In the ovaries, we identified 412 unique DEGs among the four behavioral groups and 132 unique DEGs among these four groups in the brains. The number of DEGs between each comparison is found in table S2. The full differential expression results are found in the supplemental material. Overall, comparison of isolated subordinates and natural guards revealed little to no differences in gene expression (1 DEG in the ovaries and 0 DEGs in the brain), in contrast to isolated subordinates versus subordinates from an active social nest (73 DEGs in the ovaries and 94 DEGs in the brain). These brain and ovary DEGs showed highly significant overlap with DEGs independently identified between natural guard and subordinate individuals in the brains (49/88 shared DEGs, p<0.001) and ovaries (41/68 shared DEGs, p<0.001) from Saleh and Ramírez, 2019. Hierarchical clustering based on expression patterns of the 412 ovary DEGs revealed two clusters mostly corresponding to reproductive (dominants and subordinates) and nonreproductive (isolated guards and natural guards) phenotypes (Fig. 4). This gene set is enriched for multiple GO-terms, including “signal” and “transmembrane” (full results Table S4). Furthermore, this gene set includes genes known to be associated with reproductive and social behavior in insects; for example, DNMT3, broad-complex, corazonin receptor, yellow-g, yellow-g2 (Drapeau et al., 2006; Paul et al., 2006; Okada et al., 2016; Gospocic et al., 2017). In the brains, Hierarchical clustering of the 132 brain DEGs clustered subordinates separately from the three other behaviors (Fig. 5). This gene set was enriched for multiple GO-terms including “signal” and “vision”. This gene set also includes genes known to be involved in insect social and reproductive behavior, such as hexamerin 70c, hormone receptor-like 38, prohormone 2, prohormone 3 (Okada et al., 2016; Shpigler et al., 2019).

**Figure 4.**
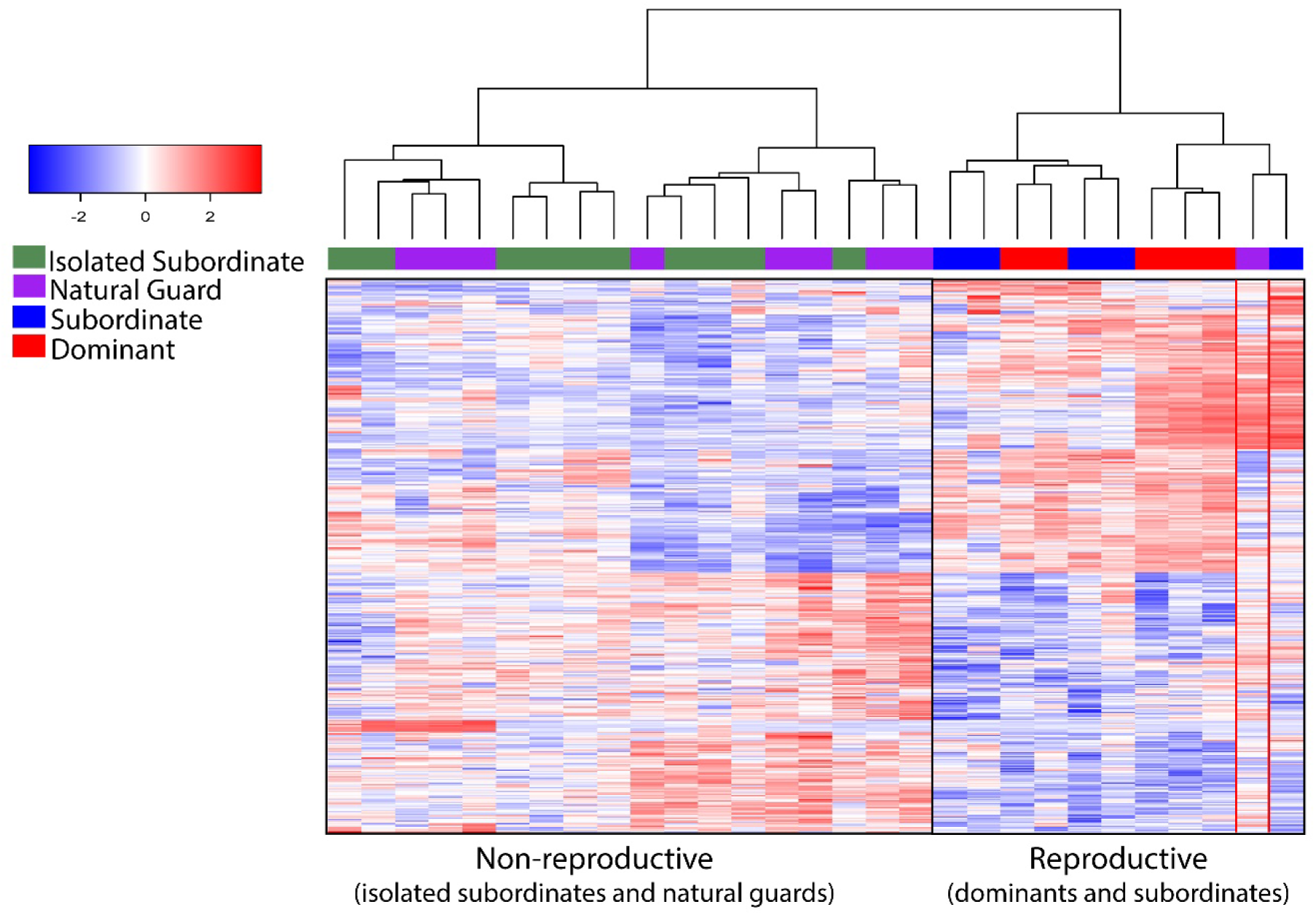
Hierarchical clustering of samples based on expression of 412 pairwise DEGs identified among behaviors in the ovaries. Color key shows the log_2_ scaled expression relative to the mean value for each gene. Nonreproductive and reproductive clusters are highlighted with black boxes, with one natural guard that does not cluster according to reproductive state is highlighted in red. Sample clustering was based on using Euclidean distance with the Ward.D2 clustering method.

**Figure 5.**
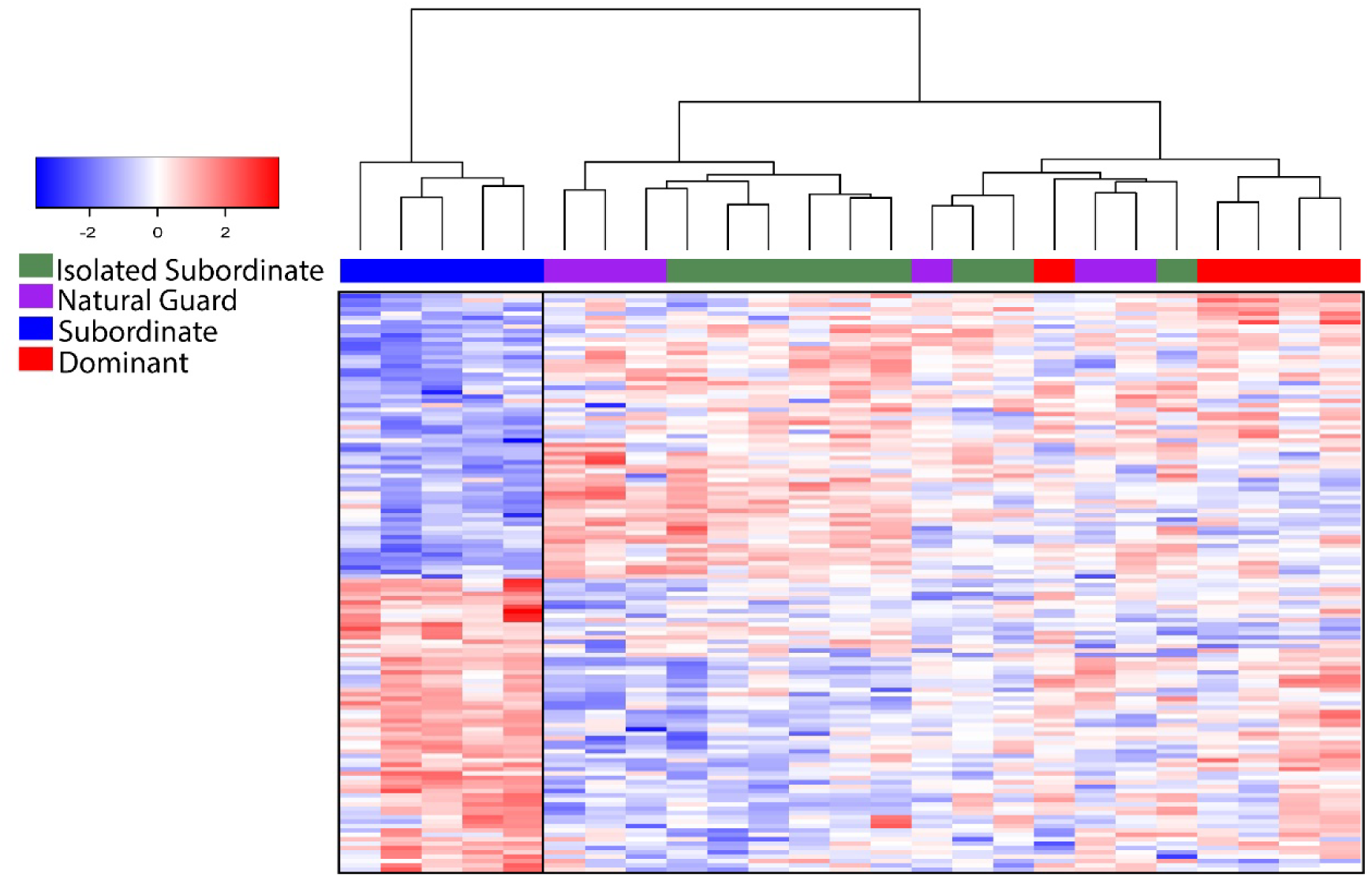
Hierarchical clustering of samples based on expression of the 132 pairwise DEGs identified among behaviors in the brain. Black boxes highlight the subordinate cluster and the cluster containing the other three behaviors. Color key shows the log_2_ scaled expression relative to the mean value for each gene. Sample clustering was based on using Euclidean distance with the Ward.D2 clustering method.

We also performed specific contrasts in the EdgeR model to compare sampled behaviors based on reproductive state and initial dispersal strategy. For reproductive state we performed a contrast between the two reproductive groups (dominants/subordinates) and the two nonreproductive groups (isolated subordinates/natural guards). For initial dispersal strategy we compared females that emerged in the nest and stayed as helpers (subordinate/isolated subordinate) and females that dispersed to begin their own nest on emergence (dominants/natural guards). Considering the reproductive state contrast, we find 514 DEGs in the ovaries and 109 DEGs in the brains. In the ovaries, 318 genes are shared between this reproductive contrast and the 412 DEGs identified among all pairwise behavioral comparisons above (p<0.001). This is in line with hierarchical clustering supporting reproductive state as the major factor driving differential expression patterns among behavioral groups in the ovaries. In the brains, 64 of the 109 genes are shared with the 132 pairwise DEGs identified across behaviors, a greater overlap than expected by chance (p<0.001), though clustering in the brain by behavioral DEGs does not correspond primarily to reproductive states. Considering differing dispersal strategies (subordinates and isolated subordinates vs dominants and natural guards), we find less genes overall, with 9 DEGs in the brain and 3 DEGs in the ovaries.

In summary, our differential expression analysis finds that isolated subordinates and natural guards show similar expression profiles at identified DEGs (as seen in hierarchical clustering) and we find only 1 DEG across tissues between the two behaviors. In addition, reproductive changes rather than initial dispersal strategy appear to be associated with more expression differences in both the brains and ovaries.

### Gene network analysis identifies modules of genes that are highly correlated with ovary size

Using WGCNA, we identified 13 modules of co-expressed genes in the brains and 13 modules in the ovaries and examined correlations between these modules and ovary size. Multiple module eigengenes, which represent the first principal component summarizing the expression of genes within each module (Langfelder & Horvath, 2008), were correlated with ovary size changes across the sampled brain and ovary tissues. The full results and analysis are detailed in supplemental tables and the accompanying R code, including identities of the genes present in each module (table S5), connectedness values (kME) for genes in these modules (tables S6 and S7), gene enrichment analysis for the focal modules discussed below (table S4), and a correlation matrix comparing all the modules in both tissues with ovary size index (Fig S7). Several especially strong connections between co-expression modules and ovary size are highlighted here. Following FDR correction for 26 comparisons to ovary size, eigengenes from 8 ovary modules and 2 brain modules were significantly correlated with ovary size variation. Two ovary module eigengenes showed especially strong correlation with ovary size. Ovary module six, which consisted of 371 genes, showed a strong negative correlation with ovary size (r = - 0.89, p<0.001, Fig. 6) and ovary module 10, consisting of 116 genes, showed a strong positive correlation with ovary size (r = 0.89, p<0.001, Fig. S8). The genes in ovary module 6 were significantly enriched for the “DNA replication” KEGG pathway, while there were no significantly enriched terms for module 10. In the brain, module 3, a large module of 1,256 genes, showed the highest correlation (r = -0.54, p= 0.028, Fig S9) with ovary size variation. This module was enriched for several terms, including “coiled coil,” “transducer,” and “nucleic acid binding,” and included multiple genes with known associations with social behavior, such as syntaxin-1A and dopamine receptor D1 (Kocher et al., 2018; Sasaki, 2010). This module was also correlated with ovary module 6 (r = 0.62, p<0.01), suggesting co-expression across tissues. In line with this, 71 of the 350 possible overlapping genes in ovary module 6 are shared with brain module 3, a larger overlap than expected by chance (p<0.001).

**Figure 6.**
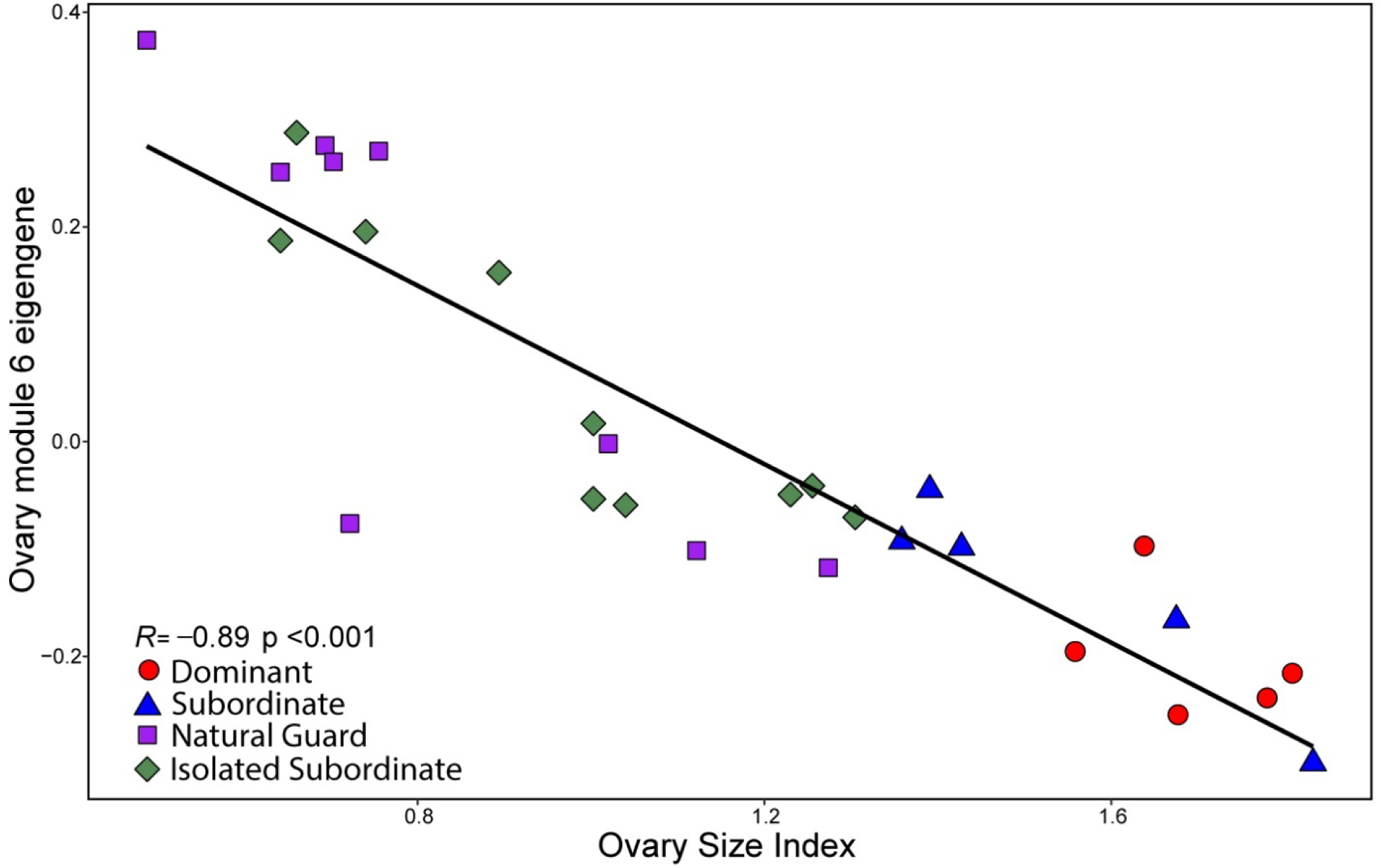
Correlation between eigengene values and ovary size index measurements from module 6, containing 371 genes detected with WGCNA of ovary transcriptome data. Spearman correlation coefficient is shown. The P-value was FDR adjusted for 26 ovary size comparisons.

### Cross-study gene list comparisons

We compared genes identified through differential expression and WGCNA to those associated with worker related reproductive plasticity in *A. mellifera* and *M. genalis*, two species showing complex and simple eusocial organization, respectively. We focus these comparisons on genes related to the reproductive differences we identify, to test the hypothesis that genes involved in reproductive plasticity identified here overlap with genes associated with reproductive plasticity in other species of eusocial bees. In the ovaries, genes upregulated in natural guards and isolated subordinates (nonreproductive) versus dominants and subordinates (reproductive) significantly overlapped with genes upregulated in both *M. genalis* worker versus queen abdomens (64/126 shared genes, p<0.001) and in the abdomens of non-egg laying vs egg laying honey bee workers (35/196 shared genes, p<0.001). Considering the opposite contrast, genes upregulated in reproductive vs nonreproductive *E. dilemma* individuals, these genes significantly overlap with genes upregulated in *M. genalis* queens versus workers (65/163 shared genes, p = 0.044) but not with egg laying versus non-egg laying workers (36/227 shared genes, p = 0.24).

We also compared the gene lists from *M. genalis* and *A. mellifera* to the genes identified in the two WGCNA modules with the strongest associations with ovary size (ovary modules 6 and 10). Genes present in module 6 significantly overlapped with genes differentially expressed between *M. genalis* queens and worker abdomens (159/227 shared genes, p<0.01) and *A. mellifera* non-egg laying vs laying workers (89/266 shared genes, p<0.001). In contrast, module 10 did not significantly overlap with *M. genalis* queen versus worker abdomen DEGS (49/80 shared genes, p = 0.5) or *A. mellifera* non-egg laying vs laying honey bee workers (26/92 shared genes, p = 0.06).

## Discussion

In this study we disrupt the social structure of small colonies to assess the behavioral and reproductive plasticity of *E. dilemma* subordinate females, finding that socially isolated subordinate bees are highly flexibly, capable of expressing largely the same behavioral, physiological, chemical, and gene expression changes that dispersing foundress bees show naturally across solitary behavioral phases. In addition, using gene network and differential expression analysis, we identify sets of genes strongly associated with reproductive plasticity that overlap with genes associated with worker physiology in bees exhibiting eusocial behavior. This suggests that the lack of nonreproductive workers in *E. dilemma* is not due to a lack of physiological plasticity in subordinates, which show high behavioral and physiological flexibility.

### The initial effect of social disruption on *E. dilemma* subordinates

Our experimental manipulation of 14 individuals resulted in 10 subordinate individuals that transitioned to guard behavior and were collected, although there was variation in the timing of this transition (Table S1). Three of the 14 individuals disappeared or died before transitioning to guarding behavior and one individual died after transitioning to guarding behavior before collection. Four of the 10 individuals that transitioned to guarding behavior stopped provisioning additional brood cells after isolation and two of 10 provisioned only one additional brood cell before guarding. This suggests that some individuals may be responding to nest disruption by beginning to guard their brood early, relative to foundresses that have transitioned to guarding. This may be what is driving the slightly smaller average brood size of isolated subordinates (Fig. S4). This response could be due to stress imposed by the treatment itself. However, of the four of 14 bees that disappeared after isolation but before collection, three of them finished provisioning at least one brood cell after isolation. Only one bee disappeared the day following isolation without continuing to forage, which suggests that the isolation treatment may not have been an extreme source of stress to most individuals. Alternatively, it could be that the changing social environment affects decisions about optimal nest defense and brood size, though the data presented here cannot address this. Although some individuals appeared to respond directly to isolation by quickly transitioning to guard behavior, others continued provisioning long after isolation. One individual, for instance, provisioned an additional seven brood cells following isolation, ultimately guarding a relatively large brood of nine. In general, it is unclear what mix of environmental and genetic factors influence the size of the first brood. Additional work to disentangle sources of variation on brood size is necessary to provide greater insight into behavioral mechanisms underlying the transition from foraging to guarding behavior.

### Reproductive physiology and behavior may differentially influence transcriptomic and chemical variation

Although the transition to guarding behavior is accompanied by a clear reduction in ovary size in isolated subordinates (Fig 2), reproduction is not necessarily the only influence on the phenotypes we examined. In the brain, for instance, hierarchical clustering of samples based on DEGs from pairwise comparisons of behaviors revealed two primary clusters that were not correlated with reproductive state (Fig. 5). Instead, samples grouped subordinates separately from the other three behavioral groups (dominants, natural guards, isolated subordinates). This is largely in line with findings from Saleh and Ramírez, et al., 2019, where brain variation associated with social hierarchy clustered individuals according to foraging/non-foraging behavior rather than reproductive state, such that dominants and natural guards clustered together and subordinates and foraging foundresses (not sampled in this study) clustered together. Considering this, clustering of the isolated subordinates with dominants and guards is consistent with these individuals making the same behavioral changes associated with a transition from foraging outside the nest to remaining inside the nest during the day. The “vision” GO term was enriched in the gene set, and it may relate to a transition from foraging outside the nest to remaining in the dark nesting environment. In addition, NMDS analysis of CHC profiles revealed mostly the same clustering pattern as seen in brain DEGs, with subordinates clustering separately from the other sampled behaviors, which are all non-foraging. This pattern may be explained by differential light exposure among individuals, along with behavioral and physiological variation, which can have a strong impact on CHC profiles due to UV exposure and degradation, possibly influencing these clustering patterns (Hatano et al., 2020).

In our samples, subordinate bees were younger on average than the other sampled groups (typically true in natural nests), which may additionally affect the resulting transcriptomes and CHC profiles. However, the absolute difference in age between isolated subordinates/natural guards and dominants is likely greater than the difference in age between isolated subordinates/natural guards and subordinates. Dominants sampled in this study have, for example, completed the guard phase and then participated in social nesting for multiple weeks. Consequently, if age itself is primarily driving the phenotypic variation we identify, the effect would not be strictly linear. If the patterns we observe were driven primarily by age and not foraging/non-foraging status, for example, we might expect to see natural guards, isolated subordinates, and subordinates grouping together, as these three groups should be closer together in age on average than any group is to dominant bees. Furthermore, analysis of independently collected *E. dilemma* transcriptomes from Saleh and Ramírez, 2019 (appendix 1, Figs A7 and A8), shows a signal of behavioral clustering with the DEGs identified in this study, suggesting that these DEGs are likely associated with behavior and physiology independent of any sampling biases that could be present in this dataset.

The clustering pattern we identified in CHC profiles and pairwise brain DEGs contrasts with the sample clustering that we identified based on the pairwise ovary DEGs, which revealed two clusters mostly corresponding to the reproductive/nonreproductive phenotypes also seen clearly in the ovary size index measurements (Fig. 2). Although clustering based on behavioral DEGs in the brain was not primarily based on reproductive differences, these differences did clearly influence brain transcriptomes, as seen by the subset of overlapping DEGs between the nonreproductive/reproductive contrast and the pairwise DEGs among all behaviors. Furthermore, the brain gene module most strongly correlated with ovary size variation contained a relatively large amount of genes, some of which have been previously found to be associated with social behavior in other species.

### Foundresses and subordinates have the same physiological, chemical, and transcriptomic potential

Our data strongly suggest that subordinate bees are totipotent and can express the full spectrum of reproductive and behavioral changes seen during the foundress to guard transition. Indeed, natural guards and isolated subordinates are largely indistinguishable by all phenotypes we examined. This, along with the lack of body size differences among *E. dilemma* behavioral groups, suggests that subordinate behavior is probably not the result of strong developmental differences limiting the plasticity of some individuals. Consequently, it seems unlikely that the foundress versus subordinate trajectories in *E. dilemma* are strictly determined by large nutritional differences, as these would likely be reflected in body size differences (Lawson et al., 2017). However, it is possible that subtle nutritional and/or developmental differences still underly these behaviors and require additional investigation to uncover. This contrasts with several other well-studied bees showing simple social behavior, such as some halictids and small carpenter bees, in which maternal manipulation of nutrition strongly influences the body size and social trajectory of offspring (Lawson et al., 2017; Kapheim, 2016).

This begs the question then, what determines whether a newly emerged female will disperse or stay as a subordinate? It has been recognized in several orchid bee species that early eclosing females are much more likely to remain in their natal nest and become subordinates compared to later eclosing females (Andrade-Silva and Nascimento, 2012; Augusto and Garófalo, 2009). Although this generally appears to be true in *E. dilemma*, the pattern is not always consistent (N Saleh, personal observation) and does not explain why the number of subordinates varies among nests. In addition, this observation is somewhat confounded by the fact that, because *E. dilemma* nests rarely grow beyond 1-2 subordinates, later eclosing females may have little opportunity to remain in the nest. Thus, correlates with eclosion order cannot easily be teased apart without further experimental manipulation.

In the social allodapine bee *Exoneura bicolor*, eclosion order and not developmental or nutritional factors is the proximate determinant of dominance status, so post-eclosion hierarchy determination is known to occur in bees (Schwarz and Woods, 1994). Furthermore, in *Euglossa townsendi* social nests, individuals can transition back and forth between dominant-like behaviors and subordinate-like behaviors, irrespective of age, such that the social hierarchy shifts over time among a group of individuals (Augusto and Garófalo, 2004). In *E. dilemma*, one possibility is that newly emerged females could undergo some decision-making process, integrating information about the local availability of nesting resources (*e.g*. pollen, resin), the current state of the nest, the age and condition of current occupants, and the likelihood of inheriting the nest, before remaining as a subordinate or dispersing as a foundress. Experimental manipulation of these factors is a necessary next step in clarifying the role of developmental versus post-eclosion factors in determining offspring trajectory.

### Why do *E. dilemma* social groups lack nonreproductive workers?

Considering the question posed by Roberts and Dodson (1967), “why, then, has there been no evolution of distinct worker and reproductive castes among these bees?” we can, from a proximate perspective, rule out the hypothesis that individuals lack the reproductive plasticity to express worker-like physiology. The data presented here show that subordinates will exhibit nonreproductive phenotypes that involve genes associated with worker physiology in eusocial species. This leaves us with the question then, if subordinate reproductive physiology can be dynamically regulated, why doesn’t this happen in social groups, leading to nonreproductive workers? Other bees, especially some Halictid bee species, such as *M. genalis*, can form small social groups comparable in size to those in *E. dilemma* that still contain nonreproductive workers (Kapheim et al., 2012). Thus, social group size itself does not have to impose a limit on the evolution of nonreproductive workers.

One possibility is that subordinate eggs are functioning as a type of trophic egg and that disrupting this process may have fitness costs on social individuals. Many stingless bee species, despite their derived form of eusociality, have workers with activated ovaries that lay trophic eggs for the queen (Wille, 1983). Furthermore, trophic egg laying appears to be ancestral in the stingless bees, though, in contrast with orchid bees, they have repeatedly evolved nonreproductive workers (Gruter, 2018). Consequently, comparative analysis of orchid bee and stingless bee oophagy behaviors may be especially useful in understanding how trophic eggs evolve and function and whether these traits influence the evolution of nonreproductive workers. Ultimately, in *E. dilemma* and generally across orchid bees, additional data is necessary to investigate the evolutionary forces (or lack thereof) that have shaped reproductive interactions in social groups.

## Conclusions

These data show that individual *E. dilemma* females exhibit a high degree of phenotypic plasticity, with each female capable of large behavioral and reproductive changes regardless of their initial foundress/subordinate trajectory. Furthermore, these reproductive changes involve genes associated with worker physiology in eusocial species, suggesting that *E. dilemma* subordinates are capable of worker-like nonreproductive physiology even though this physiology and social behavior appear to have evolved unlinked. As such, *E. dilemma* represents a unique case in the corbiculate bees where functional reproductive division of labor has evolved via behavioral (oophagy) but not physiological specialization. Future study of the specific cues triggering guarding behavior and nonreproductive physiology in *E. dilemma* may provide insight into the mechanisms enabling and/or preventing the evolution of nonreproductive workers during social evolution.

## Acknowledgements

We thank the members of the Ramírez lab for their help and advice throughout the project. We thank Nikki Hochberg, Ron Phenix, and all the staff at the Fern Forest Nature Center for their support of this project. We thank Joe Patt and Aleena Tarshish Moreno for their help. Funding was provided to S.R.R. by the David and Lucile Packard Foundation and the National Science Foundation (DEB-1457753). Funding was provided to N.W.S. by the Daphne and Ted Pengelley Award (University of California, Davis) and the National Science Foundation (PRFB award number: 2109456). Funding was provided to J.H. by the Studienstiftung des deutschen Volkes.

## Author contributions

N.W.S. and S.R.R. designed the project. N.W.S and J.H. collected samples. N.W.S. performed sample processing and data analysis. All authors participated in writing and revising the manuscript.

## Data Availability

Code and data files to reproduce the analysis in the manuscript have been deposited in Dryad to be published on acceptance at https://doi.org/10.25338/B8TK96. Sequencing data can be found at NCBI Bioproject accession PRJNA750777.

## Notes

### Competing Interest Statement

The authors have declared no competing interest.

